# Analysis of histone antibody specificity directly in sequencing data using siQ-ChIP

**DOI:** 10.1101/2023.03.08.531745

**Authors:** Ariana Kupai, Robert M. Vaughan, Scott B. Rothbart, Bradley M. Dickson

## Abstract

We previously developed sans spike-in quantitative chromatin immunoprecipitation sequencing (siQ-ChIP), a technique that introduces an absolute quantitative scale to ChIP-seq data without reliance on spike-in normalization approaches. The physical model of siQ-ChIP predicted that the IP step of ChIP would produce a classical binding isotherm when antibody or epitope was titrated. Here, we define experimental conditions in which this titration is observable for antibodies that recognize modified states of histone proteins. We show that minimally sequenced points along an isotherm can reveal differential binding specificities that are associated with on- and off-target epitope interactions. This work demonstrates that the interpretation of histone post-translational modification distribution from ChIP-seq data has a dependence on antibody concentration. Collectively, these studies introduce a simplified and reproducible experimental method to generate quantitative ChIP-seq data without spike-in normalization and demonstrate that histone antibody specificity can be analyzed directly in ChIP-seq experiments.

## INTRODUCTION

Chromatin immunoprecipitation sequencing (ChIP-seq) is the gold standard technique for mapping the distribution of histone post-translational modifications (PTMs) on the genome^1, 2^. The biophysical theory of ChIP-seq shows that, if performed in a properly controlled manner, the method can also be a quantitative technique, where the abundance of immunoprecipitated (IP’d) species at a defined genomic interval can be measured^3, 4^. Efforts to establish a quantitative scale for ChIP-seq have relied on the addition of exogenous chromatin standards prior to IP^5–7^. The “spiked-in” reagents serve as a normalizer across experiments, and their use is purported to be necessary for quantification of sequencing data^8^. Spike-ins are often used to measure relative gains/losses in a PTM between two samples after a perturbation^5^. Because of the relative scale, conditions of spike-in experiments must be closely matched to compare datasets. Unfortunately, most published ChIP-seq data lack sufficient reporting of key parameters (i.e., chromatin input concentration, immunoprecipitated DNA mass, volumes, etc.) to critically evaluate the scale and reproducibility of the data^9^. Additionally, spike-in methods are often time consuming, laborious, and introduce new sources of potential variation in the experimental workflow. An absolute, quantitative scale for ChIP-seq without the use of spike-ins simplifies comparisons of quantitative ChIP-seq data between labs and institutions, thus providing immeasurably greater benefit and information to the community per ChIP-seq experiment than is currently garnered.

We previously developed a physical and quantitative scale with which to interpret ChIP-seq outcomes^3, 10^. This method, sans spike-in quantitative ChIP-seq (siQ-ChIP), is an inherently quantitative approach that does not require spike-ins for signal normalization. The physical model of siQ-ChIP predicts that the IP step of ChIP-seq is a competitive binding reaction that produces an isotherm of mass capture as a function of antibody or epitope concentration. Here, we defined experimental conditions that capture these titration dynamics, and we investigated whether sequencing points of mass capture along the isotherm could change the composition of IP’d DNA.

Our studies confirm a central hypothesis derived from the siQ-ChIP theory^3^; sequencing points along a binding isotherm results in differential peak responses for antibodies that recognize multiple epitopes. Through the action of this differential peak response, siQ-ChIP can distinguish strong and weak antibody-epitope interactions. Strong (high affinity) interactions typically belong to epitopes that the antibody is selected against and would be considered on-target interactions. On the contrary, weak (low affinity) interactions may be called off-target interactions. Different scenarios of antibody:chromatin binding are possible. An antibody may have one observable binding constant, meaning it interacts with the same affinity to all epitopes. In this situation, technologies like histone peptide microarrays^11^ can help inform on whether the antibody is solely recognizing the epitope of interest or is recognizing multiple epitopes with the same affinity, the former being the best for a ChIP-seq experiment and the latter being the worst. Both scenarios would represent an antibody with a narrow spectrum of binding constants. Likewise, an antibody could display a broad spectrum of binding constants, where an antibody could bind most strongly to the intended target epitope but also exhibit weaker binding to other epitopes. We sequenced the IP material from multiple histone post-translational modification (PTM) antibodies and observed both narrow and broad spectrum antibodies. Throughout this manuscript, we refer to these two classes of antibodies as “narrow” and “broad” as opposed to “on-” and “off-target”, to reflect the spectrum of binding constants within the antibody:chromatin interaction.

This work highlights the significance of experimental parameters, such as antibody concentration, in the interpretation of for ChIP-seq data. Notably, this distinction of narrow versus broad histone PTM antibody binding spectrum can be determined by sequencing at a low depth (12.5M reads per IP), making characterization of antibody specificity within a ChIP-seq experiment inexpensive and feasible without the need for spike-in reagents.

## RESULTS

### Development of an Optimized siQ-ChIP Protocol

We first sought to streamline and optimize experimental conditions necessary to generate titratable ChIP binding isotherms compatible with the siQ-ChIP analysis pipeline. This optimized protocol, presented in step format in the methods section, also removes unjustified complexity from other quantitative ChIP protocols^6, 12^. The protocol requires optimization of micrococcal nuclease (MNase) digestion of DNA to mono-nucleosome fragments and generation of a chromatin concentration standard (**Fig. 1A**). Once optimized, the ChIP workflow from cells in culture to isolated DNA fragments takes only 1.5 days with approximately 4 hours of hands-on time (**Fig. 1B**). **Figure 1** contains data showing successful execution of these steps. These data serve as benchmarks to be met before moving on to sequencing.

**Figure 1.**
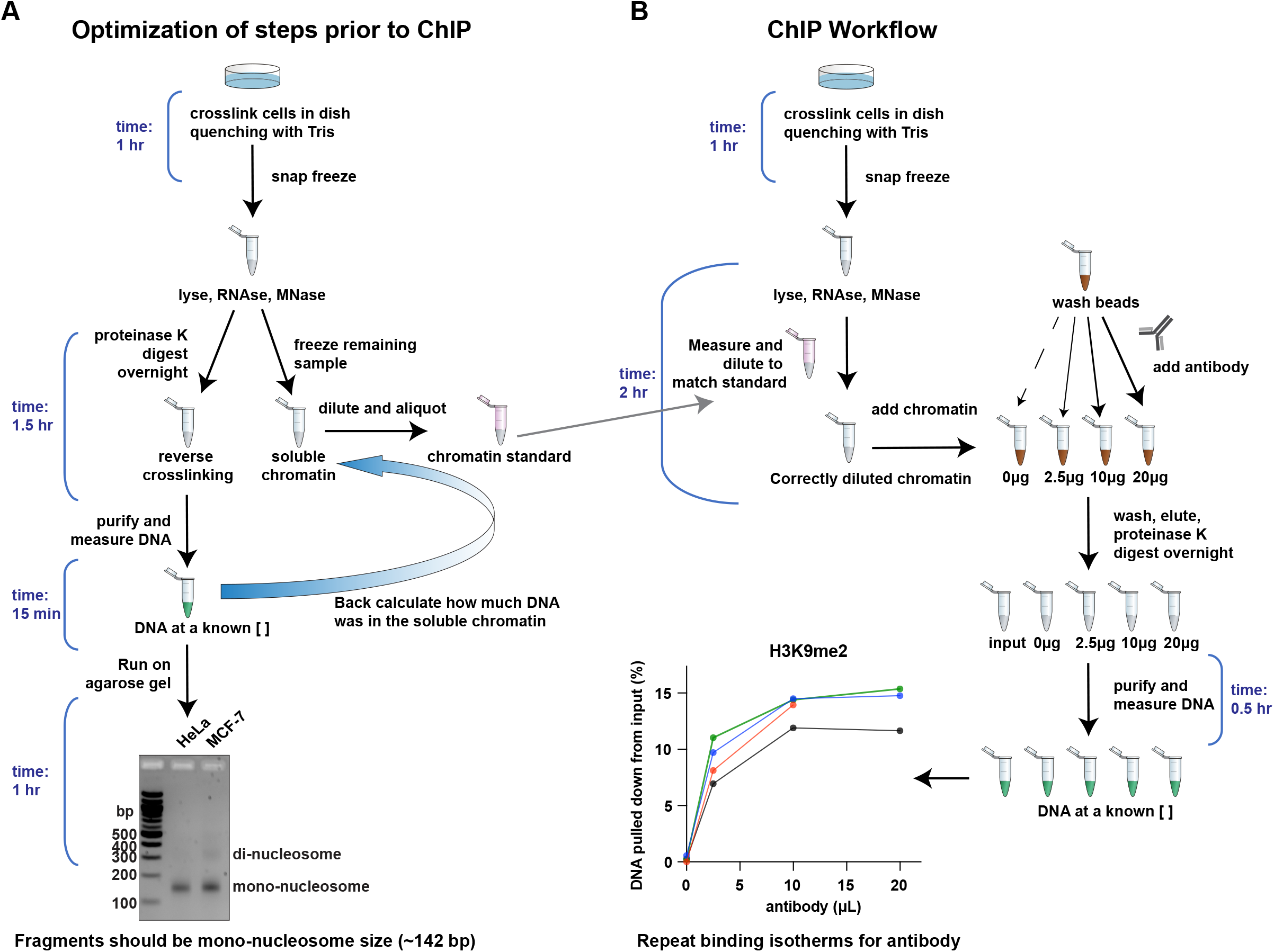
Sample optimization and enrichment workflow for siQ-ChIP (**A**) Optimization of steps prior to siQ-ChIP. These steps should be completed before an IP experiment. UV-imaged DNA gel analysis of MNase digestions from two cell lines shows enrichment for mononucleosome-sized chromatin fragments. Chromatin was incubated with 75 U MNase for 5 min at 37 °C. The gel is a 2.5% agarose gel with 50 ng of DNA from HeLa and MCF-7 cells. (**B**) Stepwise illustration of the siQ-ChIP experimental workflow. Using H3K9me2 from Fig. 3B as an example. a binding isotherm is measured, where increasing antibody causes an increase of IP’d DNA, until saturation is reached. The amount of IP’d DNA and the shape of the isotherm is similar across replicates.

Proper control over critical protocol steps contributes to increased reproducibility of ChIP experiments. We evaluated several of these steps in the process of optimizing this protocol. Formaldehyde crosslinking has been used to stabilize DNA-protein interactions for decades^13^. After crosslinking, formaldehyde needs to be quenched to limit reactivity. In ChIP protocols, quenching is often performed using glycine. It has been noted however, that glycine is unable to form a terminal product with formaldehyde^14^, casting doubt on its ability to effectively stop crosslinking^15^. We used the protocol described herein to ChIP for lysine 18 acetylation on histone H3 (H3K18ac) using either 125 mM glycine or 750 mM Tris as a formaldehyde quencher. Notably, both quenching methods IP’d the same amount of DNA (**Fig. 2A-B**) and had overlapping percentages of DNA captured from input (**Fig. 2C**). The MNase digestion of fragment lengths was also the same between both quenching methods (**Supplementary Fig. 1A**). Typically, 2.5 M glycine is added directly to formaldehyde to achieve a final concentration of 125 mM glycine^16–18^. Using this method of quenching resulted in more variable mass of IP’d DNA (**Supplementary Fig. 1B**) but did not affect MNase activity (**Supplementary Fig. 1C**). We therefore recommend removing formaldehyde prior to the addition of quenching reagent. Across biological replicates in all quenching conditions, an observable binding isotherm was obtained; more antibody resulted in more IP’d DNA until saturation was reached (**Fig. 2A-C**). Isotherms increased in amount of DNA IP’d until reaching saturation when titrating antibody, consistent with the IP step being a competitive binding reaction^3^. We moved forward with Tris quenching in our protocol due to the concerns of reproducibility described above.

**Figure 2.**
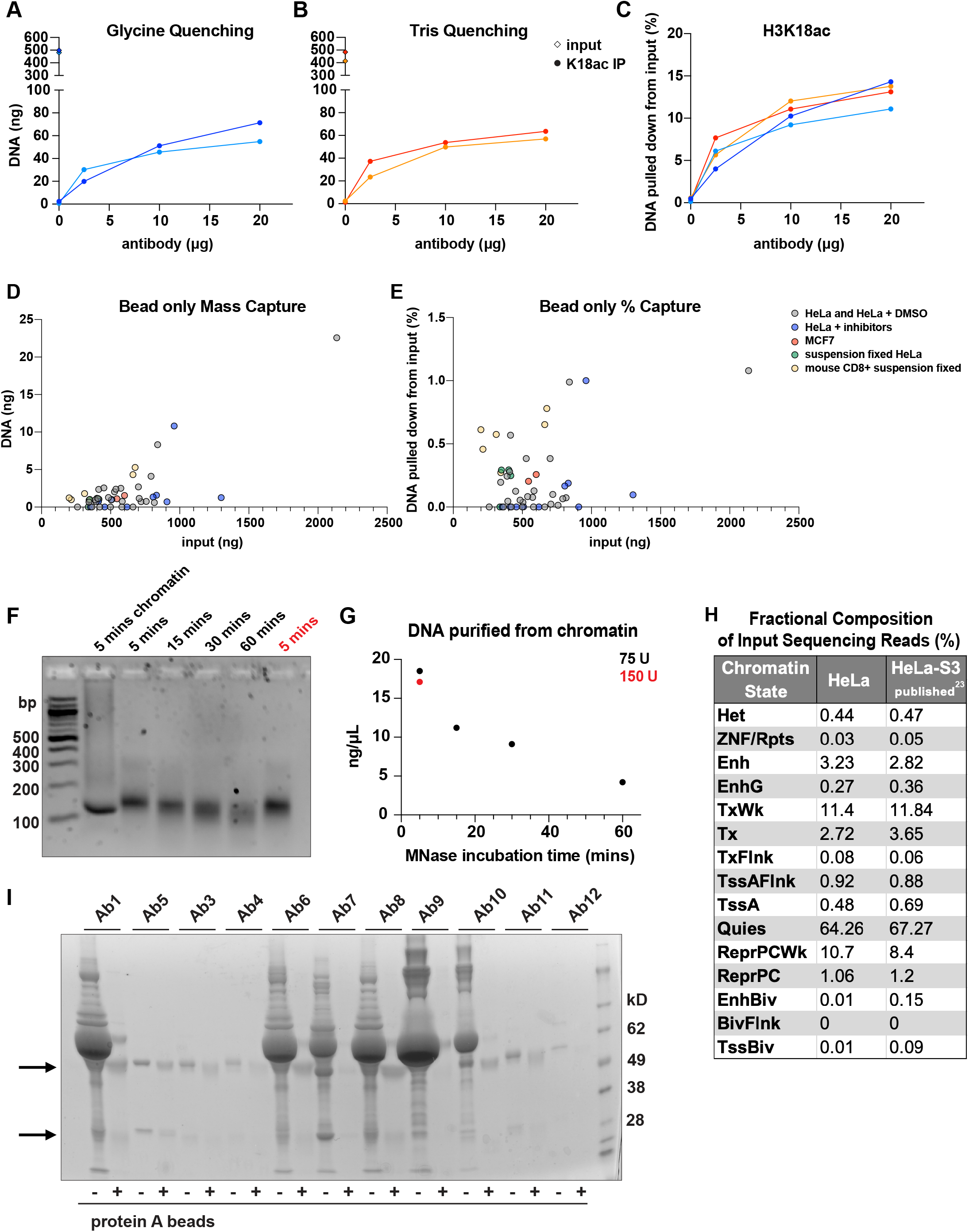
Critical siQ-ChIP protocol steps H3K18ac antibody (Ab11) titration in HeLa cells after removing formaldehyde and quenching with either (**A**) 125 mM glycine or (**B**) 750 mM Tris reported as mass of DNA captured or (**C**) percent IP’d from input. Repeat isotherms were generated from cells crosslinked at the same time but IP’d on different days. Protein A bead only capture from Tris quenching experiments as (**D**) mass or (**E**) percent IP’d from input. (**F**) 2.5% agarose gel of hTERT RPE-1 cells treated with the indicated amounts of MNase for the indicated times at 37 °C. Lane 1 is unpurified chromatin; all other lanes are 100 ng of column-purified DNA. (**G**) Mass concentration of DNA purified from hTERT RPE-1 cells from panel C. (**H**) Fractional composition of HeLa input reads across the 15-state ChromHMM model compared to published HeLa-S3 values^23^ (see also Supp. Fig. S6e, average percentage values used when HeLa-S3 values were not greater or less than one standard deviation away from the average). (**I**) Protein A bead elution after antibody incubation.

For a reproducible isotherm, bead only capture of DNA needs to be minimized. We note that our optimized protocol does not include bead pre-clearing or blocking steps common to many methods. We measured increased bead-only DNA capture with increased concentration of input chromatin (**Fig. 2D**). When plotting bead-only DNA mass as a percentage of input DNA, capture never exceeded 1.2% across almost 50 replicates from different cell types and treatments (**Fig. 2E**). Therefore, bead pre-clearing and blocking steps were deemed unnecessary. Removing these steps resulted in a simplified protocol that was free of non-specific bead-chromatin interactions. In our own practice, bead-only capture greater than ∼1.5% of input disqualifies samples from sequencing (or further IP).

There are two common ways to fragment DNA in ChIP experiments: sonication and MNase digestion. Even with optimization, sonication produces fragments in a range of sizes from 100- 800 bp^19^. MNase however, is an endo-exonuclease that nicks DNA and continues to cleave nucleotides until reaching a nucleosome, resulting in mono-nucleosome-sized DNA fragments^20^. The reproducibility of a small range of fragment sizes makes MNase a superior way to fragment chromatin for downstream quantitative purposes. In sequencing, the average DNA fragment length is used for scaling, and quantification will be less accurate for samples displaying a complex distribution of fragment sizes, as is the case for sonication. Additionally, using MNase over a sonicator makes this method more accessible to labs that do not have access to specialized equipment.

To determine optimal MNase digestion conditions, several MNase concentrations and incubation times were tested (**Fig. 2F-G**). Notably, MNase-treated chromatin was not always representative of purified DNA fragments. Chromatin incubated with 75 U of MNase for 5 min (**Fig. 2F**, lane 1), appeared to contain a large smear of DNA followed by a prominent mono- nucleosome-sized fragment. When DNA was purified from the MNase digested chromatin (**Fig. 2F**, lane 2), the smear was gone, and an additional di-nucleosome band was seen. For this reason, it is recommended to only use purified DNA to check MNase digestion. We also tested the effects of MNase incubation time. Over time, the di-nucleosome band seen using 75 U of MNase for 5 min disappeared, and the mono-nucleosome band shifted to a lower weight and became smeared (**Fig. 2F**). The smearing and decreased mass of the mono-nucleosome band represented over-digestion, which was also evident by the decreased amount of DNA recovered after longer MNase incubations (**Fig. 2G**). Increasing the amount of MNase from 75 U to 150 U did not change fragment size or DNA amount recovered (**Fig. 2F-G**). These experiments informed the use of digestion conditions of 75 U for 5 min per 10 cm dish of HeLa cells at 80% confluence. These digestion conditions were sufficient for three different cell types (HeLa, MCF7, and primary mouse CD8+ T cells) (**Fig. 1A, 2F**), suggesting MNase fragmentation of DNA does not need to be optimized for each sample type, and rather should be reproducible based on similar input chromatin amounts. We have sequenced input chromatin obtained from MNase digestions several times at 100 million reads and see no evidence of biased recovery of the genome, consistent with a previous report^21^. To demonstrate this, we plotted the fractional composition of Mnase digested HeLa input reads binned in a 15-state chromatin annotation model of the genome (ChromHMM)^22^. The percentage of input reads in each annotation from Mnase-digested HeLa cells was consistent with published sonicated HeLa-S3 input values from the ENCODE project^23^ (**Fig. 2H**).

An often-unreported step of ChIP is determination of compatibility between the antibody and beads. To test antibody-to-bead binding, a panel of histone PTM antibodies typically used in ChIP was incubated with Protein A magnetic beads, and bound material was eluted (**Fig. 2I**). The heavy and light chain of an antibody are 50 kD and 25 kD, respectively (**Fig. 2I, arrows**). It is clear which antibodies were not affinity purified due to the large number of bands of additional molecular weights. The large band at ∼62 kD is likely BSA in the storage buffer. This experiment can be used to troubleshoot an inability to IP DNA. For example, Ab7 did not IP any DNA in a ChIP experiment (not shown), and this antibody did not bind to magnetic beads (**Fig. 2I, lanes 11-12**). Antibody identities are listed in **Supplementary Table 1**.

To match chromatin concentration across all ChIP experiments, we use a chromatin standard. A chromatin standard is saved, frozen chromatin from a previous experiment where a small fraction underwent DNA purification and quantification, thus providing a source of chromatin of known DNA concentration. A chromatin standard needs to be measured repeatedly when using the Qubit Fluorometer because of the fluctuating value of Qubit standards, which are used to calibrate the instrument. The general workflow of how to make the chromatin standard is outlined in **Fig. 1A** and detailed in protocol steps 4-5. We found the 2^nd^ Qubit standard, which determines what relative fluorescent unit (RFU) value signifies 10 ng/μL of DNA, had a wide range of values. The value of the 2^nd^ Qubit standard directly impacted the measured concentration of the chromatin standard (**Supplementary Fig. 2**). To narrow the variation in measurements caused by the Qubit, chromatin and DNA were only measured when the 2^nd^ Qubit standard measured greater than 7000 RFU (**Supplementary Fig. 2, red arrow**). In our experience, when the 2^nd^ Qubit standard was less than 7000 RFU, the 2^nd^ Qubit standard needs to be better mixed before addition to Qubit buffer/dye mixture or a new 2^nd^ Qubit standard needs to be obtained. Our chromatin standard was from HeLa cells, but when applied to MCF-7 cells, still enabled accurate dilutions, suggesting that the standard chromatin measurement could be applied even if taken from a different cell type.

### Generation of Reproducible Antibody Titration Isotherms

An antibody titration is used to determine the chromatin-antibody binding isotherm. The isotherm is a landmark for reproducibility, giving others a well-defined target for repeating past work. In addition, the chromatin-antibody isotherm verifies the expected outcome for a competitive binding reaction, confirming a ChIP-seq experiment displays observable sensitivity to reaction conditions. The binding isotherms from four IPs of lysine 9 di-methylation on histone H3 (H3K9me2) are shown as DNA mass (**Fig. 3A**) and percent DNA captured from input (**Fig. 3B**). All measurements are reported in **Supplementary Table 1**. The amount of H3K9me2 IP’d was reproducible across two different cell types, with a maximum variance of a 1.5-fold change across experiments (**Fig. 3A**). In all experiments, the shape of the isotherm was the same, with mass of DNA IP’d increasing until 10 μg of antibody, when saturation was reached. We sequenced the input DNA, and DNA from the 2.5 μg and 10 μg IPs, from both HeLa replicates (**Fig. 3C**). The y-axis in our browser tracks is the siQ-ChIP capture efficiency, which is equal to the ratio of the IP reads over the input reads for a certain genomic region, multiplied by the proportionality constant α (see Eq. 6 of Reference 10). α is a globally applied constant that maintains proportionality between the IP’d material and the sequencing reads^3^. siQ-ChIP browser tracks are normalized on a scale of 0 to 1, with 1 representing 100% of input capture. As a side note, because of the reference to input in all IP tracks, we do not show input-only tracks. In our four IP datasets, the peak shapes were reproducible, and the peak heights between the 2.5 μg and 10 μg IPs were reproducible. The overall apparent peak locations are similar to an H3K9me2 IP fold change over control ENCODE dataset^24^ (**Fig. 3C, green**). Of note, the ENCODE dataset is not quantitative, and the y values are relative within that single experiment. Since the y value is not related to a physical measurement, any range can be selected to view the ENCODE data. Two ranges are shown: 0-2 (top green track) and 0-6 (bottom green track) (**Fig. 3C**). The 0-6 range makes it appear that the ENCODE data and the siQ-ChIP data have the same amounts of H3K9me2 captured, but such comparisons are arbitrary because the ENCODE data does not have an absolute scale. Browser shot images of ChIP-seq data have little utility for quantitative comparisons because they can be skewed to visually suggest either similarities or differences. As described below, we suggest primarily analyzing sequencing data by using the entirety of the genome to visualize the quantitative distribution of IP reads and peaks using functional annotations of chromatin.

**Figure 3.**
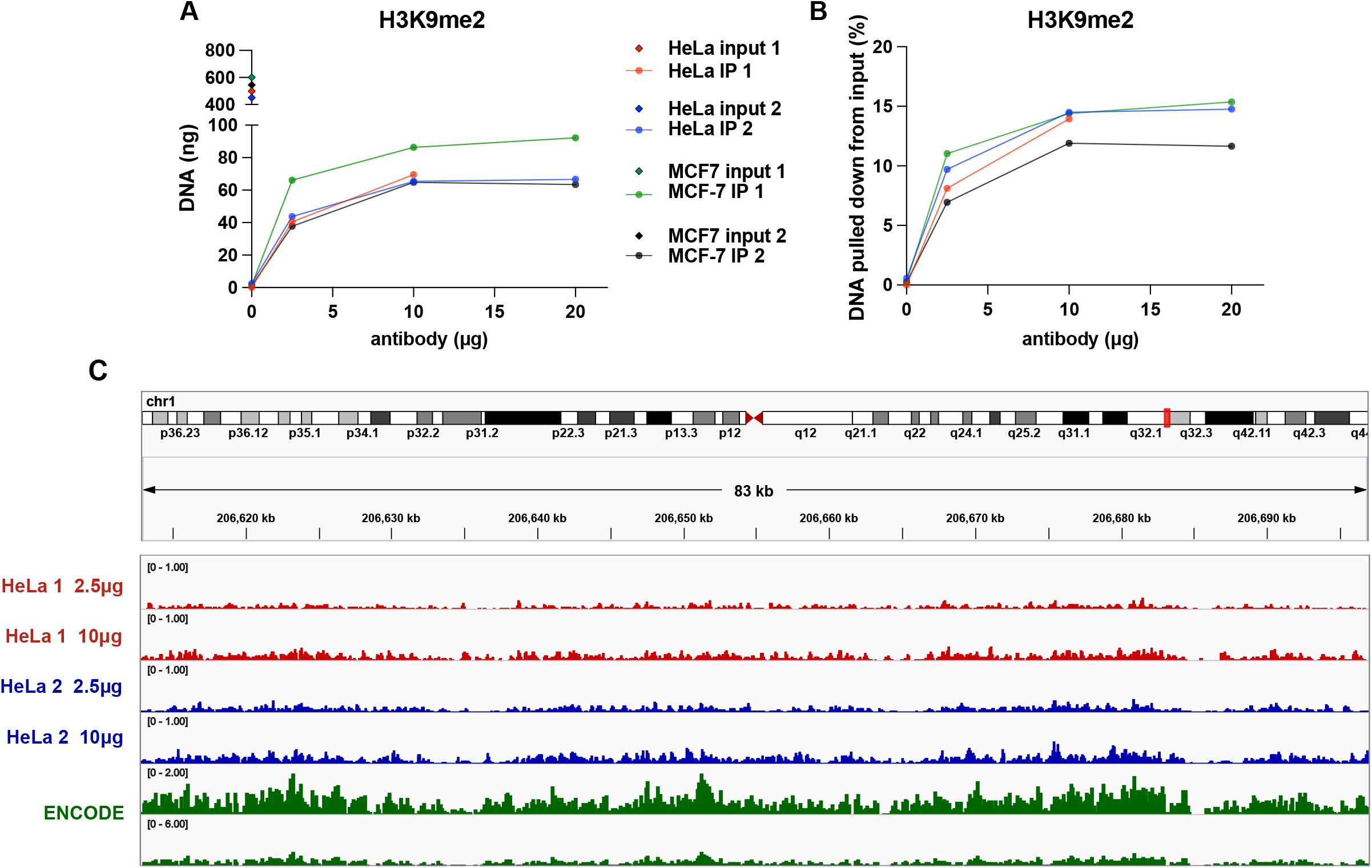
H3K9me2 DNA binding isotherm and sequencing H3K9me2 antibody (Ab5) titration in HeLa cells and MCF-7 cells displayed as either (**A**) mass IP’d or (**B**) percent of DNA IP’d from input. Repeat isotherms were generated from cells crosslinked at the same time but IP’d on different days. (**C**) Genome browser window of hg38 chr1:206612859-206697494 for Ab5 sequencing at 2.5 μg and 10 μg IPs from both HeLa replicates. ENCODE H3K9me2 IP fold enrichment from file ENCFF645RCC, experiment ENCSR328AVV shown for comparison at ranges 0-2 (upper) and 0-6 (lower). IGV Version 2.8.9.

DNA mass from IP’d material is not typically reported for ChIP-seq data. Omitting this information severely limits the comparisons one can make between datasets, directly contributing to the lack of reproducibility and necessity for the development of ChIP-seq data normalization approaches like spike-ins. Using the protocol described herein, one can achieve a reproducible binding isotherm for a given antibody without spike-ins and obtain an absolute scale for sequencing quantification. We have gone on to use several different antibodies in ChIP-seq in this paper and others^10^ and have yet to fail in generating a binding isotherm, unless the antibody did not bind to magnetic beads, as was the case for Ab7 mentioned above. Sensitivity of the chromatin-antibody isotherm is essential for concordance with the competitive binding model. The binding isotherms of histone PTMs saturate at different percentages of IP’d/input DNA according to PTM abundance. In our experience, immediate saturation of the chromatin-antibody isotherm at low amounts of antibody (≤ 2.5 ug for histone PTMs) for a histone PTM of at least moderate abundance (saturation occurs at ≥ 5% of input DNA) likely indicates a technical problem such as incomplete crosslinking or insufficient chromatin fragmentation.

### Global Analysis of Antibody Titration Sequencing Reads

Next, we determined how sequencing different points along an antibody titration isotherm affect the composition of IP’d DNA. We performed ChIP-seq with two commercial antibodies (Ab1 and Ab2) that recognize heterochromatin-associated lysine 9 tri-methylation on histone H3 (H3K9me3) and two commercial antibodies (Ab3 and Ab4) that recognize euchromatin-associated lysine 27 acetylation on histone H3 (H3K27ac). Antibodies were selected based on their histone peptide microarray profiles from our prior work^11^. We chose antibodies for each PTM that had either a narrow (Ab1 and Ab3) or broad (Ab2 and Ab4) binding spectrum. Three biological replicates of IP’d DNA are shown for each antibody (**Fig. 4A**). The H3K9me3 antibodies did not have a concentration reported on the product sheet, so the amount of antibody used was based on volume. The H3K27ac antibodies had a concentration reported, so the amount used was based on mass. These units are reflected in the Figures, but for ease of reading, we will use mass units in the text. Mass of DNA IP’d and input concentrations are in **Supplementary Table 2**. Ab1 and Ab3 saturated immediately at 2.5 μg of antibody and IP’d about 2% of input DNA. Since the abundances of these PTMs were very low, we believed immediate saturation in the binding isotherm at 2.5 μg was a possible outcome within the competitive binding framework of the IP and did not signify a technical problem. However, Ab2 and Ab4 did not reach saturation, even at 20 μg of antibody. Along the antibody titration, Ab2 and Ab4 IP’d about 3 – 14% of input DNA. H3K27ac and H3K9me3 are lowly abundant histone PTMs^25^. Therefore, the absence of saturation in the isotherms for Ab2 and Ab4 lead us to speculate these antibodies may bind to other epitopes.

**Figure 4.**
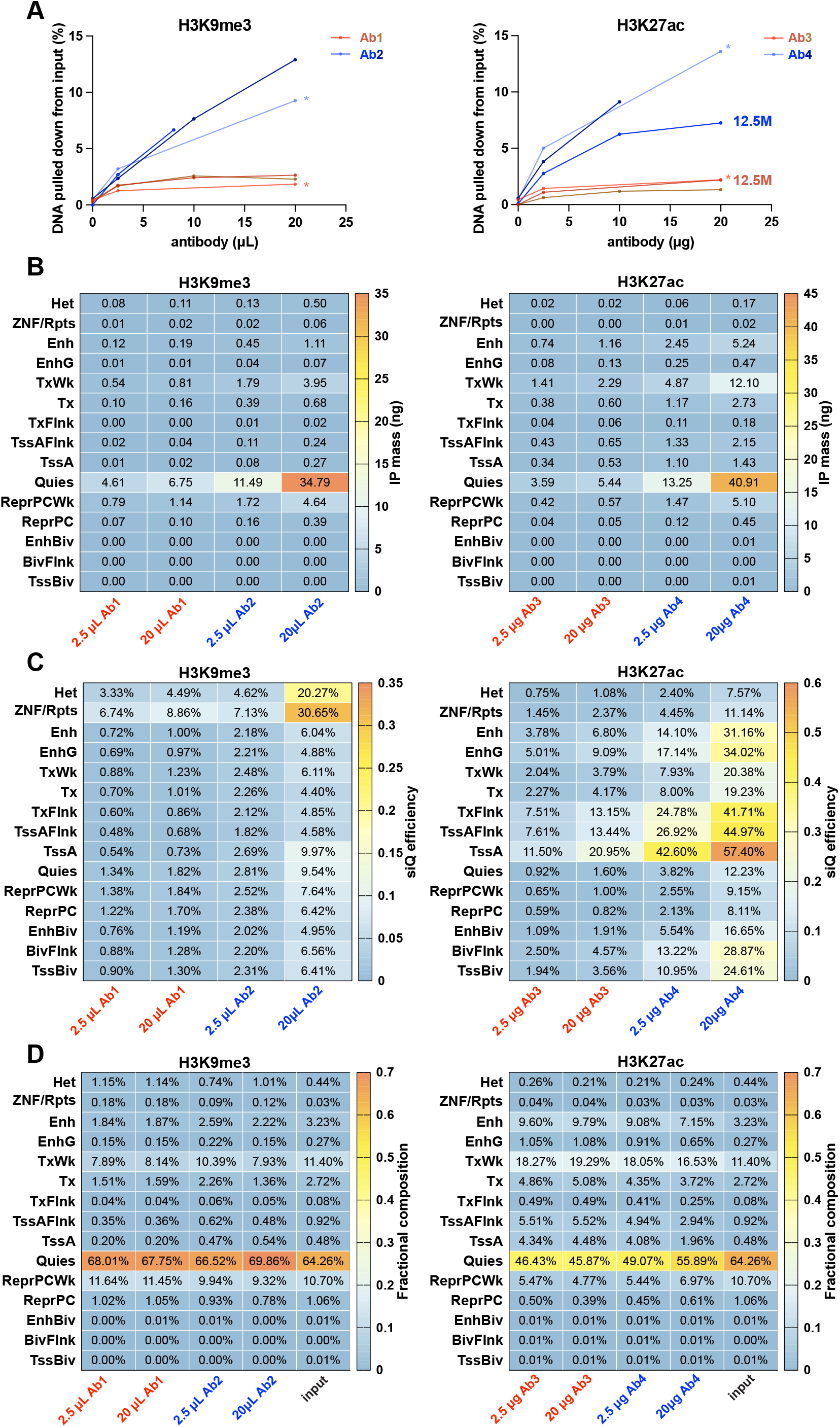
Binding isotherms and global sequencing analysis of H3K9me3 and H3K27ac antibody titration IPs (**A**) Titration of Abs1-4 in HeLa cells from three biological replicates. Data is displayed as percent of DNA IP’d from input. Replicates sequenced at 50M reads are labeled with an asterisk. Replicates sequenced at 12.5M reads are labeled with “12.5M” (see also Supp. Fig. 4). Heatmaps of whole genome analyses of sequenced reads are shown as (**B**) IP mass distribution, (**C**) siQ-ChIP capture efficiency, and (**D**) fractional composition of total reads. Genome annotations are from the 15-state ChromHMM model described in reference 22.

We sequenced the 2.5 μg and 20 μg IPs from each antibody at 50M reads. The replicates chosen for sequencing came from the same input chromatin and were denoted with an asterisk (**Fig. 4A**). Library preparation details are found in **Supplementary Table 2**. For analysis, we used the 15-state chromatin annotation model (ChromHMM)^22^ of the genome as defined by the NIH Roadmap Epigenomics Consortium^23^. We visualized the distribution of reads in these genomic states in three ways: IP mass distribution (**Fig. 4B**), siQ-ChIP capture efficiency (IP enrichment) (**Fig. 4C**), and fractional composition of total reads (**Fig. 4D**). The conceptual and mathematical derivation of these analyses is described in our prior work^10^.

Antibodies with narrow binding spectrums, (Ab1 and Ab3), had similar results at 2.5 μg and 20 μg for all analyses. Ab1 had the highest siQ-ChIP efficiency in ZNF/Rpts, and Ab3 had the highest siQ-ChIP efficiencies in TssA/flanking (transcription start sites of actively transcribed genes) and Enhancers (**Fig. 4C**). These results are consistent with on-target capture: the annotation of ZNF/Rpts is defined by H3K9me3 enrichment as determined from ENCODE datasets^26^, and TssA and Enhancers are known to be enriched for H3K27ac^27^ by use of multiple manufacturers’ antibodies^28, 29^. The efficiencies of all chromatin categories at 2.5 μg can be multiplied by a scalar value (∼1.3 for H3K9me3 and ∼1.8 for H3K27ac) to produce the efficiencies at 20 μg. Therefore, the addition of excess Ab1 and Ab3 IP’d slightly more DNA (consistent with bulk IP mass measurements in **Fig. 4A**) but did not change the distribution of IP’d DNA fragments.

Contrasting the results of Ab1 and Ab3, antibodies with broad binding spectrums (Ab2 and Ab4) showed dissimilar results in sequencing data from chromatin enrichment with two different antibody concentrations. Chromatin states with the highest siQ-ChIP efficiency were consistent within on-target antibody binding regions (**Fig. 4C**). However, gains in siQ-ChIP efficiency across all chromatin states was not uniform for Ab2 and Ab4 as it was for the two narrow spectrum antibodies. From 2.5 μg to 20 μg, Ab2 efficiencies increased from 1.9 - 4.4-fold, and Ab4 efficiencies increased from 1.3 - 3.2-fold. In conclusion, titration of Ab2 and Ab4 resulted in more DNA IP’d and also changed distributions of IP’d DNA fragments – behavior that was not observed with narrow spectrum Ab1 and Ab3.

### Peak Analysis of Antibody Titration Sequencing Reads

To compare responses between titration points for each antibody from **Figure 4**, we next analyzed peak areas of siQ-ChIP normalized tracks^10^. Recalling from earlier, the y-axis is the siQ-ChIP capture efficiency, which is normalized so 1 represents 100% of IP capture of input. In **Figure 5**, x axes (response (r)) are ratios of the peak areas in high antibody concentration IP over low antibody concentration IP. The larger the response, the larger the reduction of peak area in the low antibody data. The responses were again broken down into the 15-state ChromHMM model of the genome. If the response seen was only due to the scaling of tracks by the global proportionality constant α, then the expected response would be centered on the ratio of the high antibody’s α over the low antibody’s α. The ratio of α was ∼ 1.7 for Ab1 and Ab3 and ∼ 3 for Ab2 and Ab4 (**Fig. 5, black lines**). This means that, based on the theory of siQ-ChIP, the expectation is that peaks in Ab1/3 and Ab2/4 would be 1.7 and 3 times larger in the high antibody IP than in the low antibody IP, respectively. Peak shapes were also compared (**Supplementary Fig. 3**), but the significance of peak shape has not yet been determined^10^.

**Figure 5.**
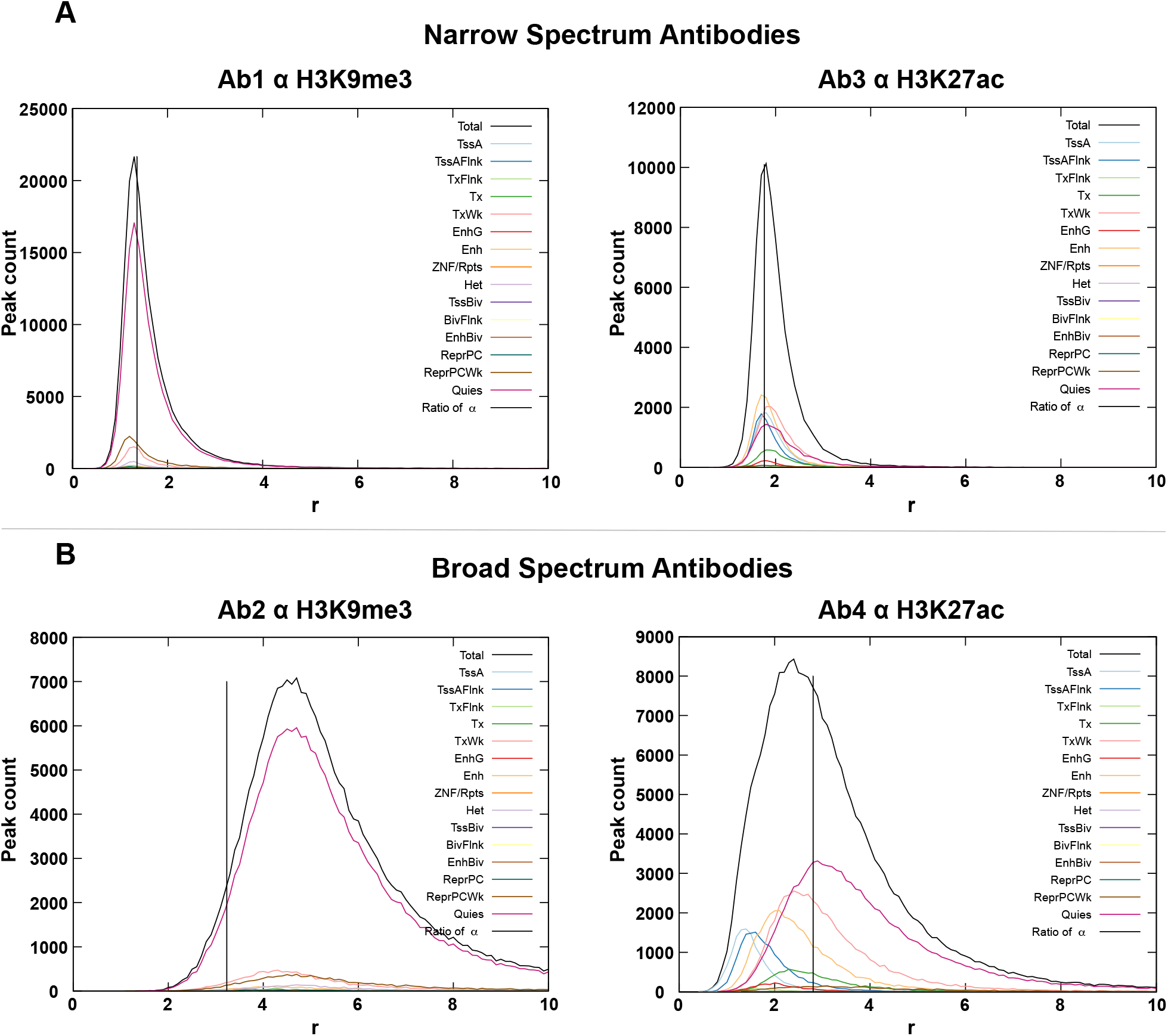
Response between titration points for H3K9me3 and H3K27ac antibodies Plots of peak response as a function of peak count across ChromHMM annotations of the genome for antibodies with (**A**) narrow and (**B**) broad binding spectrums. Vertical black line is the ratio of α in the high antibody over low antibody IP.

Antibodies with narrow binding spectrums (Ab1 and Ab3) had responses for all chromatin states centered on the ratio of α, indicating that the peak areas were only different by the ratio of scaling factors (**Fig. 5A**). On the contrary, antibodies with broad binding spectrums (Ab2 and Ab4) showed varied responses (**Fig. 5B**). Ab2 had all peak responses centered to the right of the ratio of α, and Ab4 had a wide range of responses for the different chromatin state annotations. For Ab2, the chromatin states ZNF/Rpts and Heterochromatin had high siQ-ChIP efficiency in both low and high IPs (**Fig. 4C**), but the reads may not have resulted in peaks, precluding them from response analysis. Analyzing only reads in called peaks for PTMs with lower average siQ-ChIP efficiency like H3K9me3 provides analysis of a subset of the IP data and accounts for the lack of any chromatin categories with a response less than or equal to the ratio of α. In Ab4, Quies was centered on the ratio of α, but most other categories were left shifted, with Enhancers and TssA/Flank the farthest left. Peaks in TssA/Flank regions were only 1.6-fold larger in the high antibody IP compared to the low antibody IP.

For Ab4, how was it possible that the observed response for TssA/Flank regions were smaller than the ratio of α? If we look at the fractional composition of reads, we can see the distribution of the 50M reads across the different chromatin state annotations (**Fig. 4D**). Interestingly, in Ab4 the 2.5 μg IP had 4.46% of reads in TssA and the 20 μg IP had 2.43% of reads in TssA. Since there were twice as many reads in the TssA annotation in the low antibody IP, there was a smaller response than expected for peaks in TssA regions after scaling by α’s. An increase in read number in the low antibody IP must occur for any chromatin state annotation with a response less than the ratio of α. Since α is a global scaling factor, the different responses may be caused by different binding constants of Ab4 to various regions of chromatin. This is the first time that relative binding constants for different regions of chromatin have been observed for an antibody within a ChIP-seq experiment.

### Detection of Weak Antibody Interactions

We hypothesized the cause of the differential responses for Ab4 was due to recognition of targets other than H3K27ac. To test this hypothesis, we performed *in vitro* binding assays to measure the specificity of Ab3 and Ab4. On histone peptide microarrays, both antibodies had the highest signal for H3K27ac-containing peptides (**Fig. 6A, green**). The signal from Ab3 sharpy dropped for peptides that lacked H3K27ac, while the signal from Ab4 remained elevated for acetylated H3 peptides that lacked the H3K27ac epitope. A comprehensive list of signal intensities and peptide library composition from histone peptide array experiments is presented in **Supplementary Table 4**. As off-target peptides with the highest signal for both antibodies were poly-acetylated H3 peptides, we next performed florescence polarization (FP) binding assays with Ab3 or Ab4 and a florescent H3 peptide with acetylation at lysine 9, 14, and 18 (H3K9acK14acK18ac-FAM). Consistent with array results, Ab4 had a higher affinity than Ab3 for the poly-acetylated H3 peptide (lacking H3K27ac) as shown through a left-shifted isotherm (**Fig. 6B**).

**Figure 6.**
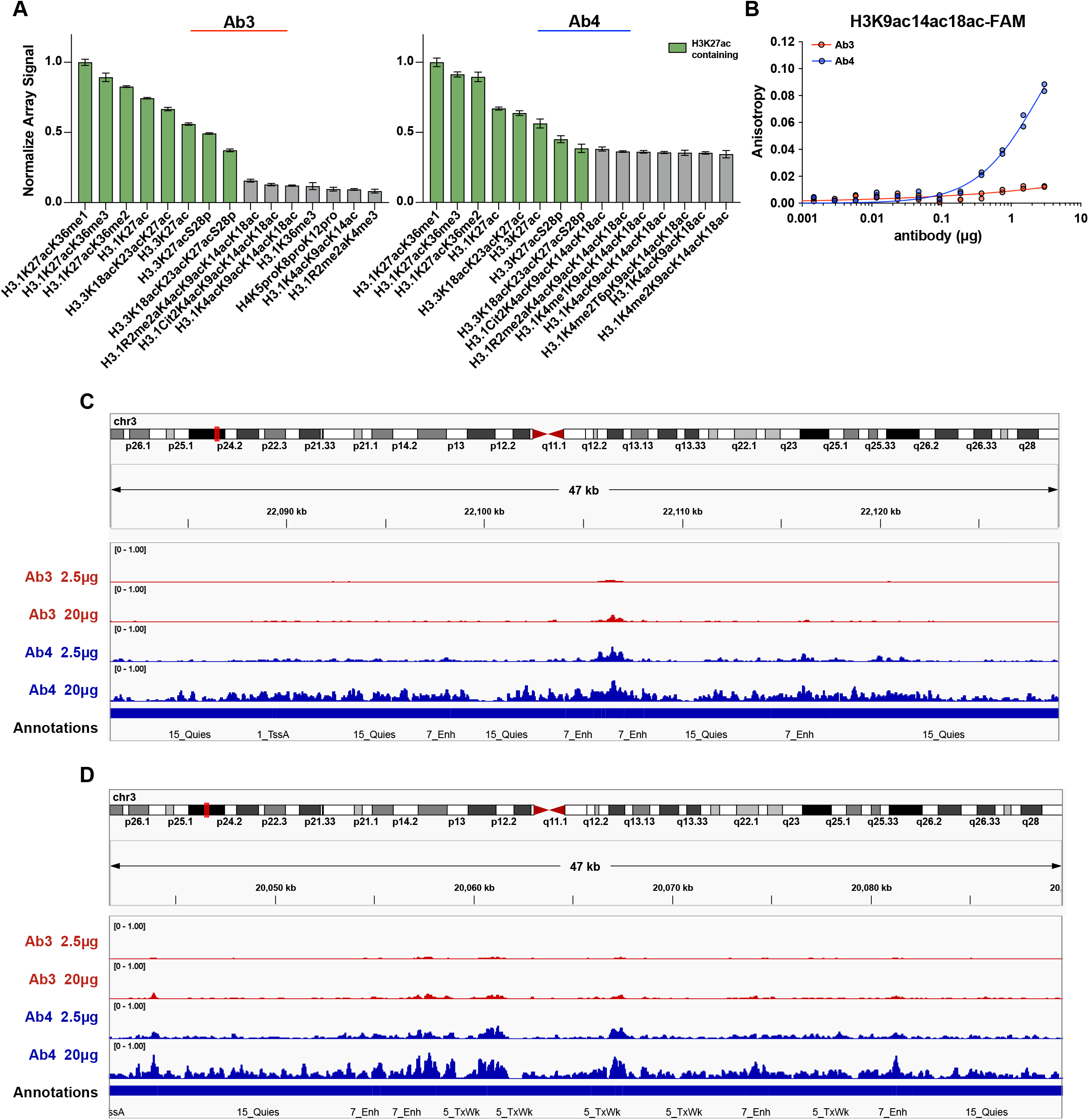
*In vitro* assessment of H3K27ac antibody specificity and distribution of sequencing reads (**A**) Intra-array normalized signal for the 15 peptides with the highest signal on histone peptide microarrays plotted as mean ± SEM from 6 technical replicates. (**B**) Fluorescence polarization binding assays of the indicated antibodies with an H3K9ac14ac18ac-FAM peptide. Technical replicates are shown. (**C**) Ab3 and Ab4 sequencing at 2.5 μg and 20 μg IPs at chr3:22081067- 22129388 and (**D**) chr3:20041655-20089976 in IGV Version 2.8.9.

We conclude that the off-target activity of Ab4 at high antibody concentrations could misconstrue PTM location. At low concentration, Ab4 binds with highest affinity to Enhancer and TssA/Flank regions (**Fig. 4C**) that are likely marked by H3K27ac. However, at high concentration, Ab4 saturates these on-target epitopes and binds to lower affinity, off-target epitopes. Instances of these additional binding events from Ab4 at high concentration can be seen in the genome browser occurring at Enhancers that have low efficiency in the Ab3 IPs (**Fig. 6C**), signifying these genomic regions may not contain H3K27ac. Additional binding also occurs at Quies and Transcriptionally Weak (TxWk) annotations (**Fig. 6C-D**). Because the capture efficiency is similar across the entire genomic window in Ab4 at 20 μg (**Fig. 6C**), it is not possible to compare peaks within one track and know what is, and what is not, target epitope. From our other work performing siQ-ChIP after p300 inhibition, we know H3K18ac is present in Enhancers, TxWk, and Quies^10^. While a low level of H3K27ac is also seen in these areas, H3K18ac is a more abundant PTM^10^, so additional Ab4 binding to these genomic regions is likely caused by recognition of off-target acetylation. As suggested by the microarray and FP results (**Fig. 6A-B**), where antibody concentration is saturating, Ab3 may be more resistant to ‘spilling over’ onto off-target epitopes. This would explain the difference in titration results and genomic distributions for these antibodies and is consistent with the theory we previously described^3^.

## DISCUSSION

This work highlights the importance of controlling ChIP-seq experimental parameters, including antibody concentration, for the accurate quantification of sequencing data. We demonstrated that different concentrations of antibody may IP more DNA and different distributions of DNA, thus changing the interpretation of histone PTM distributions across genomes. By sequencing IPs from multiple concentrations of antibody, we were able to identify which regions of DNA were likely on- or off-target, which allowed us to observe relative binding constants for an antibody towards various regions of chromatin. This work was performed with a simplified and reproducible experimental method which is included in this manuscript as a step-by-step protocol.

Consistent with our previously described theory and the treatment of ChIP as a competitive binding reaction^3^, these studies show that off-target epitope capture tracks as a function of antibody concentration. Since we have no way of determining the identity of histone PTMs on an IP’d fragment with certainty, it is critical to perform ChIP experiments under conditions of minimal antibody concentration, thus maximizing capture of high affinity (presumably on-target) interactions and minimizing capture of low affinity (presumably off-target) interactions. The practice of following a manufacturer-recommended antibody concentration in ChIP focuses on maximizing signal:background and does not account for the composition of mass capture^30^. For example, the manufacturer of Ab4 suggests to ChIP using a dilution range of 0.5-5 μg of antibody per IP but failed to mention the chromatin concentration where this dilution would be most appropriate. Instead, our studies motivate generation of binding isotherms for IP’d DNA prior to sequencing and operating below antibody saturation conditions. For antibodies that have a broad binding spectrum, precise experimental conditions can be used to observe a narrower spectrum.

We recognize that sequencing multiple points along a binding isotherm may be cost-prohibitive. To address the feasibility of antibody characterization in ChIP-seq, we also sequenced Ab3 and Ab4 low and high titration points at a depth of 12.5M reads (**Fig. 4A, replicate sequenced denoted with “12.5M”**). Even at a fourth of the depth, the same information was obtained (**Supplementary Fig. 4**) at a fraction of the cost. Testing two antibodies for a total of 50M reads on a NextSeq Mid Output flow cell with 2x75 bp paired-end sequencing reads was $410. Using the siQ-ChIP method, characterizing antibody specificity in a ChIP-seq experiment can be done inexpensively at low depths without additional consumable reagents.

We reiterate the need for reporting the following experimental measurements in ChIP experiments^3^: amount of antibody used in IP, concentration of chromatin, volume of chromatin used for IP, amount of DNA IP’d, amount of DNA taken into library preparation, amount of DNA obtained after library preparation, average fragment size after library preparation, amount of library loaded onto sequencer, and number of mapped reads sequenced. Some of these numbers are necessary to perform siQ-ChIP^10^, while others are important to address technical issues that may be apparent in sequencing results. These measurements are easily obtained by the experimentalist. We report these values for our IP and sequencing data in **Supplementary Tables 2 and 4**, and we advocate that these records should always be reported to help understand the context of the experiment and to facilitate reproducibility.

Our ChIP protocol is simple, fast, and provides clear benchmarks with which to gauge success. A limitation of this method though is the high number of cells needed. Each IP contains between one to two million cells. However, the amount of DNA for each IP is ∼500 ng, meaning our technique of solubilizing chromatin has low efficiency, as the theoretical yield is 15 μg.

Obtaining enough cells for these experiments is not problematic in our current workflow using immortalized cell lines and has not proven a limitation in processing tissue samples of at least 50 mg (not shown). Since we did not need to optimize in this direction, reducing the input mass may be possible but has not been attempted.

This workflow will be similar for all histone PTMs, but the amounts of antibody used herein for titrations are not to be used universally. We chose H3K27ac and H3K9me3 here to validate this method using PTMs associated with active and repressed chromatin states, respectively. The success of this protocol with both PTMs showed the location of the PTM in the genome does not affect or bias the IP. However, the differing abundances of these PTMs on chromatin should be considered. As determined by mass spectroscopy, H3K27ac is roughly 2% and H3K9me3 is roughly 23% of H3 peptides isolated from HeLa cells^25^. PTMs of lower abundance, such as H3K4me3 (0.4%)^25^, likely need less antibody to construct a binding isotherm before saturation. Abundance of histone PTMs from mass spectrometry can be used to generally inform about ChIP outcomes, but percentages should not be expected to be the same across techniques. Transcription factors will also likely be lowly abundant on chromatin, and we predict less antibody will need to be used in those ChIP experiments. The affinity of the antibody to the target also impacts the optimal amount of antibody to use, although this is harder to measure with certainty.

## METHODS

### ChIP Protocol

All materials can be found in **Supplementary Table 5**, ChIP Protocol Materials List. If this is your first time executing this method, you will need to complete optimization of steps prior to ChIP (**Fig. 1A**). To do this, complete steps 1-4, successfully meeting the benchmarks described. When proceeding to ChIP, make new reagents if necessary (step 1) and complete steps 2-3 again, omit step 4 and move on to steps 5-6.

#### 1. Preparation of buffers and reagents

NOTE: All buffers have a shelf life of 6 months.

1. Prepare Hypotonic Lysis Buffer composed of 20 mM Tris-HCl pH 8, 85 mM KCl, 0.5% NP- 40. Add 1 tablet of protease inhibitor per 5 mL of buffer. Buffer without protease inhibitor can be stored at room temperature. After addition of protease inhibitor, aliquot buffer and freeze at -20 °C.
2. Prepare Nuclei Lysis Buffer composed of 50 mM Tris-HCl pH 8, 150 mM NaCl, 2 mM EDTA, 1% NP-40, 0.5% sodium deoxycholate, 0.1% SDS. Add 1 tablet of protease inhibitor per 5 mL of buffer. Buffer without protease inhibitor can be stored at room temperature. After addition of protease inhibitor, aliquot buffer and freeze at -20 °C. Sodium deoxycholate is sensitive to light, so wrap the buffer bottle in aluminum foil.
3. Prepare the Binding Buffer composed of 25 mM HEPES pH 7.5, 100 mM NaCl, 0.1% NP-40. This buffer is stored at room temperature.
4. Prepare Elution Buffer composed of 25 mM HEPES pH 7.5, 100 mM NaCl, 1% SDS, and 0.1% NP-40. This buffer is stored at room temperature.
5. Prepare 1 M CaCl_2_ solution by adding 7.36 g CaCl_2_ to 42.6 mL water. This is stored at room temperature.
6. Prepare 0.5 M EDTA pH 8 solution. This is stored at room temperature.
7. Prepare 750 mM Tris pH 10-10.5 solution by adding 45.4 g Tris to 470 mL PBS. This is stored at room temperature.

#### 2. Cell fixation and collection

1. Culture cells until they reach approximately 70-80% confluency. NOTE: This is roughly 6-8 million HeLa cells. The following volumes are listed for a 10 cm dish but can be scaled by surface area for different sized cell culture vessels.
2. Rinse dish once with 10 mL D-PBS.
3. Crosslink cells for 5 min in 10 mL of 0.75% formaldehyde in D-PBS at room temperature.

1. Use a new ampule of formaldehyde for each crosslinking experiment. Dilute formaldehyde using D-PBS just prior to fixation. CAUTION: Formaldehyde is considered a hazardous chemical. Use the appropriate fume hood and safety precautions when handling.
4. Decant formaldehyde and quench cells for 5 min by addition of 10 mL of 750 mM Tris.
5. Aspirate Tris and rinse dish twice with 10 mL of D-PBS.
6. Scrape cells into 10 mL cold D-PBS, collect by centrifugation for 3 min at 300 x g, discard supernatant and snap-freeze cell pellet in liquid nitrogen.

**Pause Point:** Snap-frozen cells can be stored at -80 °C until needed.

#### 3. Chromatin isolation

1. Lyse frozen cell pellet (6 – 8 million cells) in 1 mL of Hypotonic Lysis Buffer for 30 min on ice.

1. After every volume addition/resuspension/lysis step, vortex sample for 5 s.
2. Collect nuclei (and other insoluble material) by centrifugation at 1300 x g for 5 min at 4 °C. Discard supernatant.
3. Lyse nuclei by resuspension in 150 μL of Nuclei Lysis Buffer and passage through a 27- gauge needle five times.
4. Dilute lysate by addition of 350 μL of Binding Buffer and digest RNA with addition of 5 μL of RNAse A/T1. Incubate at 37 °C for 25 min.
5. Add CaCl2 to a final concentration of 40 mM (21 μL of 1 M stock).
6. Add 75 U of MNase (3 μL of 25 U/μL). Incubate at 37 °C for 5 min.
7. Quench MNase by adding EDTA to a final concentration of 40 mM (46 μL of 0.5 M stock).
8. Dilute total volume to 1.2 mL by the addition of 625 μL of Binding Buffer.
9. Remove insoluble material by centrifugation at max speed (about 21,000 x g) at 4 °C for 5 min. Collect the supernatant containing soluble chromatin.

NOTE: If you have already made a chromatin standard, proceed to step 5.

#### 4. Generation of Chromatin Standard

NOTE: This step is only done during optimization prior to ChIP. A chromatin standard needs to be used with the Qubit Fluorometer because of the fluctuating value of Qubit standards, which are used to calibrate the instrument. The chromatin standard is used to obtain the same DNA concentration amongst all ChIP experiments.

1. Add 83 μL of Elution Buffer to 50 μL of soluble chromatin (step 3.9)

1. Freeze remaining chromatin at -20 °C.
2. Add Proteinase K to a final concentration of 15 μM (3 μL of 20 mg/mL stock).
3. Incubate at 37 °C overnight.
4. Purify DNA using the MinElute PCR Kit.

1. The final elution step should be in 30 μL of Buffer EB (10 mM Tris-Cl, pH 8.5).
5. Quantify 5 μL of the purified DNA using the Qubit dsDNA HS Assay Kit. The value of the 2^nd^ Qubit standard is important for accurate measurements (see explanation in Discussion). When measuring DNA, ensure the 2^nd^ qubit standard is within 7000-9500 RFU.
6. Check MNase digestion of DNA fragments on 1X TBE gel. MNase digestion should produce mono-nucleosome sized DNA fragments (∼150 bp).

1. Load 50-100 ng of purified DNA on a 2.5% agarose 1X TBE gel with 1X SYBR Safe.
2. Run the gel at 60V for 1-1.5 h and image under UV light. NOTE: Only continue if MNase digestion produced mostly mono-nucleosome sized DNA fragments. If mostly larger fragment sizes are obtained, see discussion for troubleshooting.
7. Use the Qubit measurement to back calculate the amount of DNA in the soluble chromatin. This is achieved using the equation:

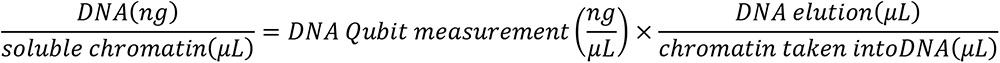

For example, if you took 50 μL of chromatin into DNA purification, eluted DNA in 30 μL, and got a Qubit reading of 4.5 ng/μL, then you would have 2.7 ng of DNA per μL of soluble chromatin:

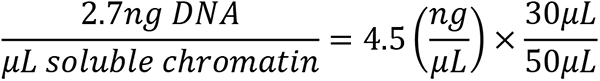
8. Use the mass concentration of DNA determined above to dilute the leftover soluble chromatin (step 4.1.1) with Binding Buffer. This is now your chromatin standard. NOTE: Our chromatin standard is at 2.88 ng DNA/μL. This concentration of chromatin has proved sufficient in ChIP experiments to immunoprecipitate a wide range of histone post-translational modifications. Optimization should be performed for targets of different abundance or scarce cell samples.
9. Aliquot the chromatin standard and freeze at -20 °C until needed. Freeze thaws do not affect the Qubit measurement. NOTE: You have now completed optimization of steps prior to ChIP. To perform ChIP, start the protocol from the beginning to obtain new soluble chromatin. Do not use frozen chromatin to ChIP.

#### 5. Standardizing chromatin concentration

1. Take 10 μL of soluble chromatin (step 3.9) and 10 μL of your chromatin standard (step 4.9) and measure them in duplicate using the Qubit (step 4.5).
2. Average the duplicate measurements for each sample. Dilute the soluble chromatin with Binding Buffer to match the measurement of the chromatin standard.
3. Save 50 μL of this chromatin for an input measurement.

#### 6. ChIP

1. Wash 25 μL of Protein A-coated magnetic beads (or whatever bead suits your antibody of interest) per IP, in batch once with Binding Buffer. For example, if you are performing four IPs, measure 4.5 x 25 μL of beads, giving yourself an additional half reaction for dead volume.

NOTE: All bead steps are performed in 1.5mL Eppendorf tubes that fit on a magnetic rack, and all centrifuge steps are done briefly in a mini centrifuge.

1. Briefly centrifuge to collect material in the lid and place tube on a magnetic rack to capture beads. Remove the liquid without disturbing the beads.
2. Resuspend beads in Binding Buffer so each sample gets 100 μL of diluted beads. In the example above, this would mean resuspending in 450 μL of Binding Buffer. Distribute 100 μL of bead mixture into each tube.
3. Titrate antibody amount by adding either 0, 2.5, 10, or 20 μg of antibody (or μL if antibody concentration is not known) to each tube of beads.
4. Bring total volume of bead+antibody mixture to 200 μL with Binding Buffer and rotate at room temperature for at least 15 min.

1. Collect bead slurry by brief centrifugation, then place on a magnetic rack until liquid is clear and remove all buffer (contains unbound antibody). NOTE: If you are performing a ChIP experiment on two samples to compare abundance of a post-translational modification, it is recommended to perform chromatin isolation and IP simultaneously to have one master mix of bead+antibody. For example, you have two cell samples and want to perform IPs using 0, 2.5, 10, and 20 μg of antibody for each sample. This is a total of 8 IPs, but we will use 9.5 x 25 μL of beads for dead volume. After washing beads, resuspend in 950 μL of binding buffer. Each bead+antibody master mix will also need dead volume, so even though we only have 2 samples, we will multiply by 2.25. We will assume this antibody is at 1 μg/μL. The volumes to use for each component are as follows:

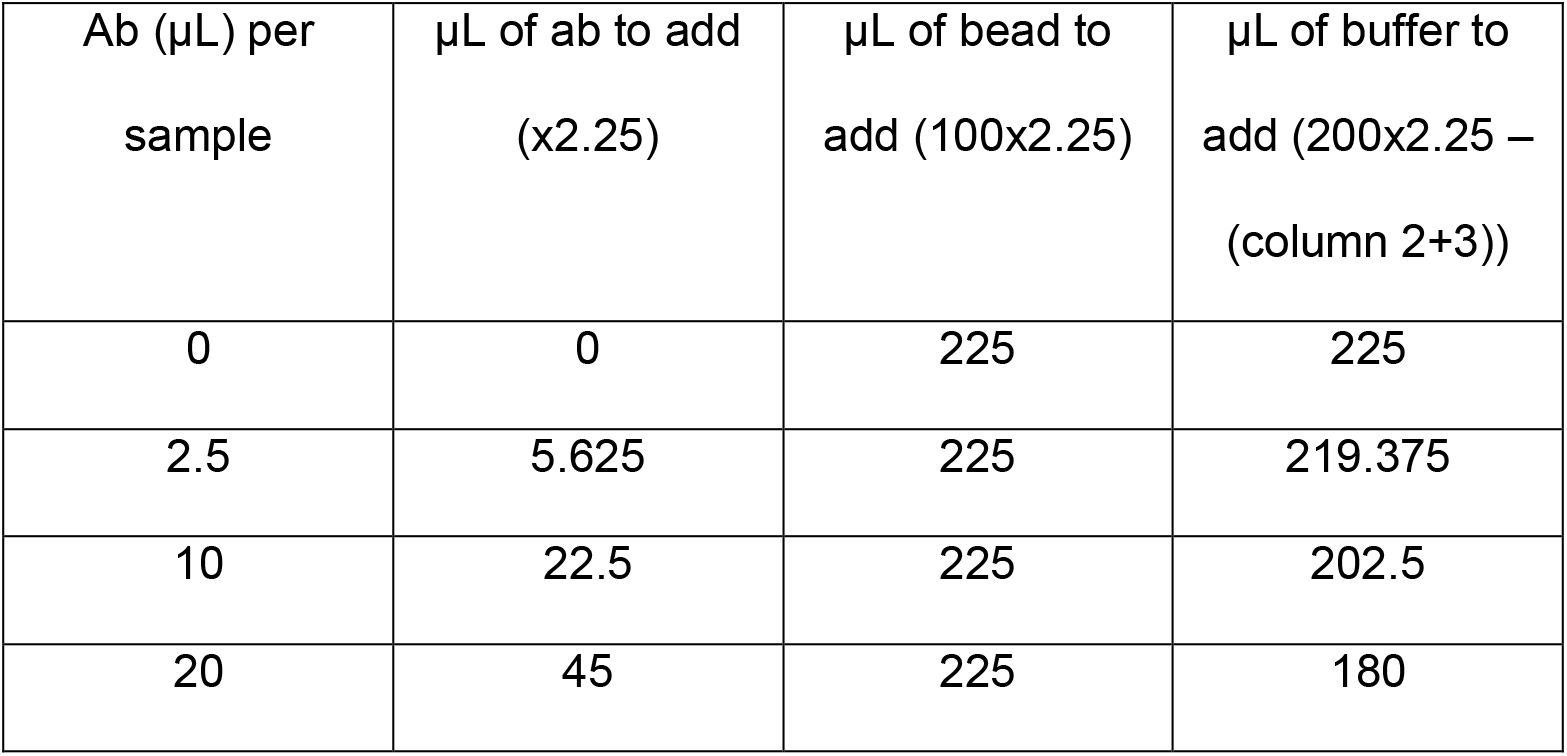 After incubation, put 200 μL from each master mix into a new tube, place tube on a magnetic rack, and remove the supernatant. Proceed to step 6.6.
5. Resuspended beads+antibody in 200 μL of soluble chromatin (step 5.2) and rotate at room temperature for at least 15 min.

1. Collect bead slurry by brief centrifugation, then place on a magnetic rack until liquid is clear and remove supernatant (contains unbound chromatin).
6. Resuspend beads in 500 μL of Binding Buffer and vortex for 10 s to wash.

1. Collect bead slurry by brief centrifugation, then place on a magnetic rack until liquid is clear and remove buffer.
7. Resuspend beads in 133 μL Elution Buffer and vortex for 10 s to elute bound material from beads.

1. Collect bead slurry by brief centrifugation, then place on a magnetic rack until liquid is clear.
2. Transfer supernatant to a new tube.
3. Bring the volume of this input chromatin (step 5.3) to 133 μL by addition of 83 μL of Elution Buffer.
8. Digest eluted chromatin (step 6.8.2) and input chromatin (step 6.8.3) by the addition of Proteinase K to a final concentration of 15 μM (3 μL of 20 mg/mL stock).

1. Incubate overnight at 37 °C.
9. Purify DNA and Quantify using the Qubit (step 4.4-4.5). The IPs plotted as percent of DNA IP’d from total available DNA (input) are given by:

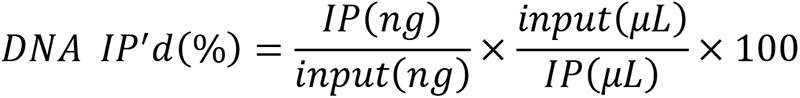
10. Check to ensure DNA fragment size is mostly ∼150 bp (step 4.6)
11. Freeze the remaining eluted DNA at -20 °C until ready for library preparation.

### Cell Culture

HeLa cells (ATCC #CCL-2) and MCF7 cells (gift from Matt Steensma) were maintained in DMEM (Gibco, 11965092) supplemented with 10% FBS (Sigma, F0926) and 1% penicillin-streptomycin (Gibco, 15140122). hTERT RPE-1 cells (ATCC# CRL-4000, Lot:70021355) were maintained in DMEM/ F12 (Gibco, 11330-032) supplemented with 10% FBS (Sigma, F0926) and 0.01 mg/ml hygromycin B (Gibco, 10687010). All cells were grown in 37°C with 5% CO2.

### Antibody capture with Protein A beads

Antibody (1 μg if known concentration, 1 μL if unknown) was incubated with 12.5 μL Protein A magnetic beads in Binding Buffer (200 μL total volume). Beads+antibody were rotated for 15 min at room temperature before being placed on a magnetic rack. Unbound supernatant was discarded. Bead-bound material was eluted with 15 μL of 1X SDS loading dye supplemented with 2-Mercaptoethanol, at 95 °C for 5 min. The beads were set on a magnetic rack, and all eluted volume was loaded onto a 4-20% gradient acrylamide gel for SDS polyacrylamide gel electrophoresis (PAGE). One μg (if known concentration, 1 μL if unknown) of antibody taken directly from the manufacturers tube was loaded onto the gel for comparison. Gels were resolved at 195 V for 1 hr followed by Coomassie staining and destaining in 10% acetic acid, 50% MeOH, and 40% H_2_O overnight.

### Sequencing and Analysis

Amount of material taken into library preparation and other information involving library concentrations, library dilutions, and sequencer loading are available in **Supplementary Table 2**. Library preparation was done using the KAPA HyperPrep Kit (Roche, KK8504) with 4 μL of Illumina adapters (IDT, UDI). Sequencing was performed using paired-end 75 bp reads on the Illumina NextSeq 500. Input libraries corresponding to H3K9me2 sequencing had > 136 M reads, and the input library corresponding to the remaining data is inputDMSO from GSE207783. IP libraries had between 48-69 M reads that passed QC, with 90% of the bases having quality scores of ≥ 30. Sequencing data were aligned to the hg38 genome as described [8]. Track building, read normalization, and other steps are described elsewhere [9]. The bw files were processed using the code on https://github.com/BradleyDickson/siQ-ChIP, updated on 11/15/2022. siQ-ChIP software automatically computes the responses using the ratio of area under overlapping peaks for compared tracks. Raw sequencing files, processed siQ-ChIP tracks, and files needed for the generation of siQ-ChIP tracks (EXPLayout and parameter files) have been deposited in NCBI’s Gene Expression Omnibus^31^ and are accessible through GEO Series accession number GSE212386. Inclusion of the files used to generate the siQ-ChIP tracks should be standard when uploading sequencing data to allow transparency and critical evaluation of data quantification.

### Histone Peptide Microarray

Histone peptide microarrays were constructed as previously described^32^. H3K27ac antibodies were diluted 1:5000 in 25 mM HEPES pH 7.5, 100 mM NaCl, 0.1% NP-40 and were hybridized to the slide at room temperature with rotation for 1 hr. Slides were washed 3 × 5 min in PBS with 0.1% Tween-20. Next, AlexaFluor 647-labeled secondary antibody (Life Technologies A- 21245, 1:5,000 dilution) was hybridized to the slide at room temperature with rotation for 1 hr. An Innopsys InnoScan 110AL microarray scanner was used to image slides, and ArrayNinja^33^ was used to quantify signals.

### Fluorescence Polarization

An H3K9ac14ac18ac peptide (amino acids 1-20) functionalized with C-terminal 5- carboxyfluorescin (FAM) was synthesized by the UNC High Throughput Peptide Synthesis Core facility. Three micrograms of each antibody were serially diluted in a black 384 well plate (Corning #3575) and 10 nM FAM peptide in 25 mM HEPES pH 7.5, 100 mM NaCl, 0.05% NP-40 were added. Polarization was measured on a Synergy Neo fluorescence plate reader (Biotek) with a 485±10 nm excitation filter and a 528±10 nm emission filter. Measurements were scaled to the lowest dilution of protein with a requested polarization of 20 milli-polarization units (mP). Anisotropy units (A) were calculated using the equation A = (2P)/(3-P).

## ACKNOWLEDGMENTS

The authors acknowledge the Van Andel Institute Genomics Core and the UNC High Throughout Peptide Synthesis Core for their services. This project was supported in part by grants from the National Institutes of Health to S.B.R. (R35GM124736) and R.M.V. (K00CA245821).

---

**Supplementary Figure 1.**
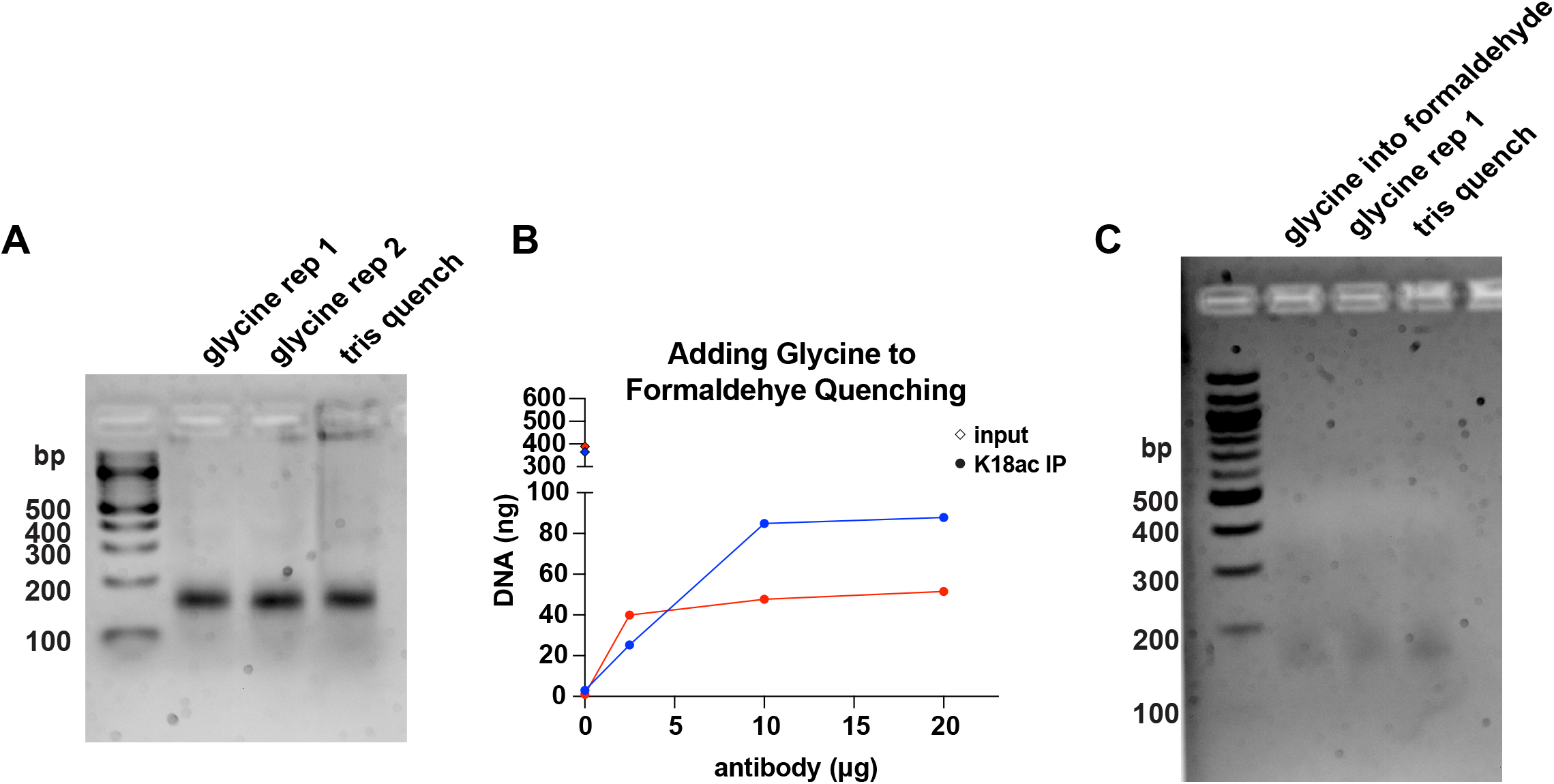
Additional optimization of the siQ-ChIP wet-bench protocol. (**A**) 2.5% agarose gel analysis of DNA from Mnase-digested input HeLa chromatin following glycine or Tris quenching. (**B**) H3K18ac antibody (Ab11) titration in HeLa cells with the direct addition of glycine to formaldehyde. (**C**) 2.5% agarose gel analysis of DNA from Mnase-digested input HeLa chromatin quenched by addition of 2.5 M glycine to formaldehyde for a final concentration of 125 mM (lane 2), removal of formaldehyde followed by 125 mM glycine (lane 3), or removal of formaldehyde followed by 750 mM Tris (lane 4).

**Supplementary Figure 2.**
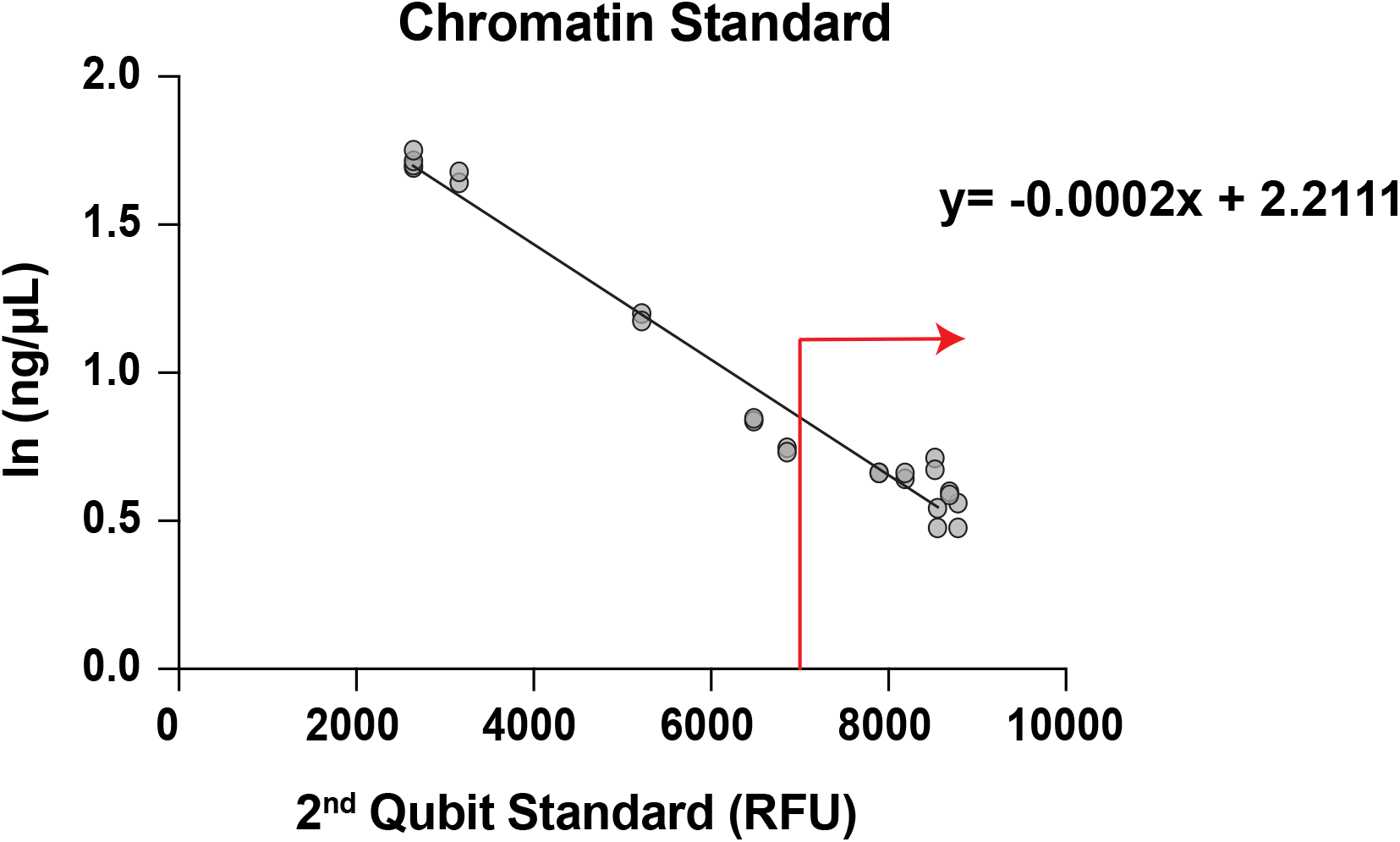
Chromatin standard fluorescence measurements across Qubit standardization concentrations Repeat measurements of the 2.88 ng/μL HeLa chromatin standard at different 2^nd^ Qubit standards (x-axis), which is used to calibrate the Qubit Fluorometer. The Y axis is the natural log of the Qubit measurement. Linear regression was plotted in excel and the best fit linear trend line is shown. RFU; relative fluorescence unit.

**Supplementary Figure 3.**
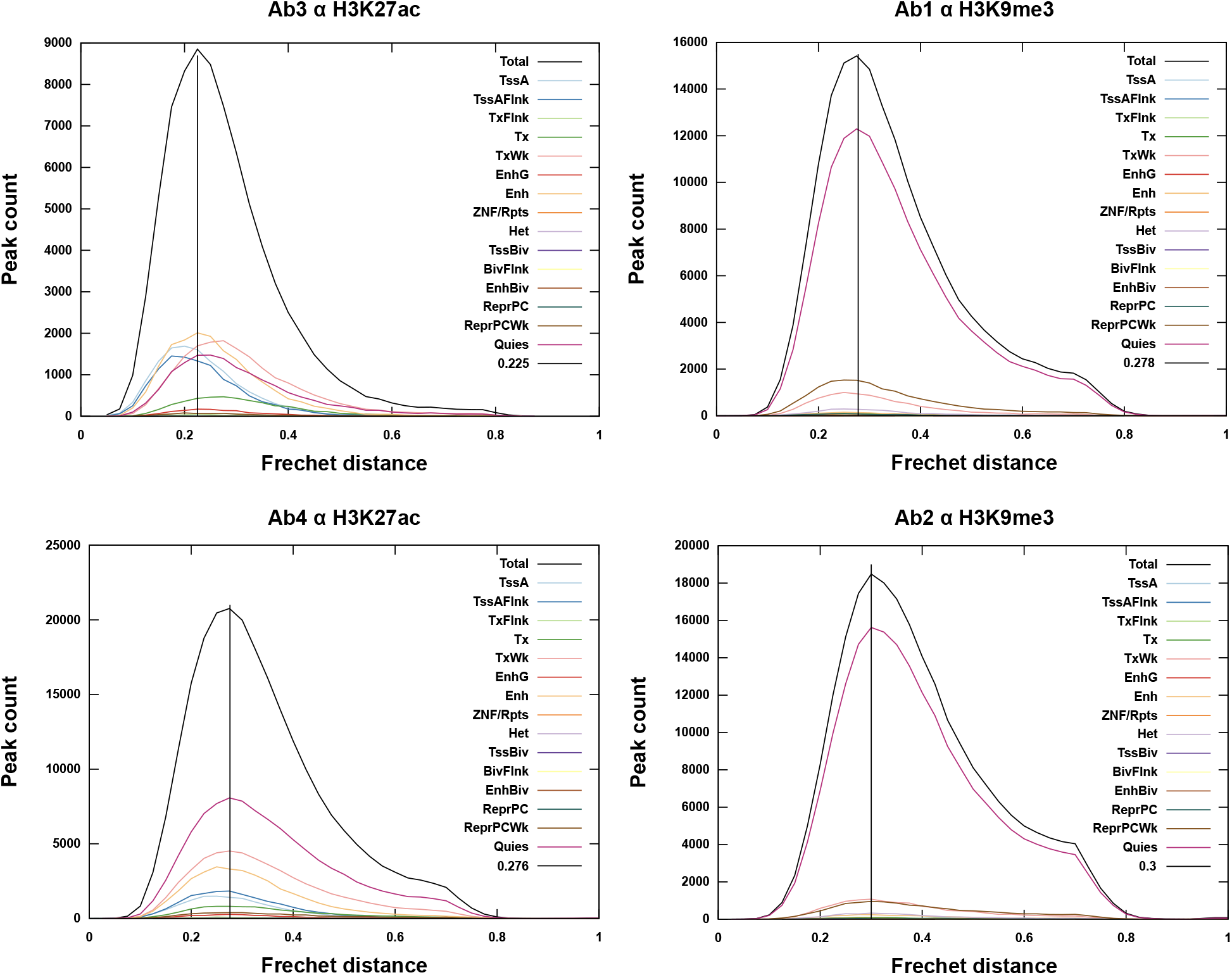
Analysis of peak shape in siQ-ChIP sequencing data Difference of peak shape between high and low antibody titrations points was calculated using the Fréchet distance as described^10^.

**Supplementary Figure 4.**
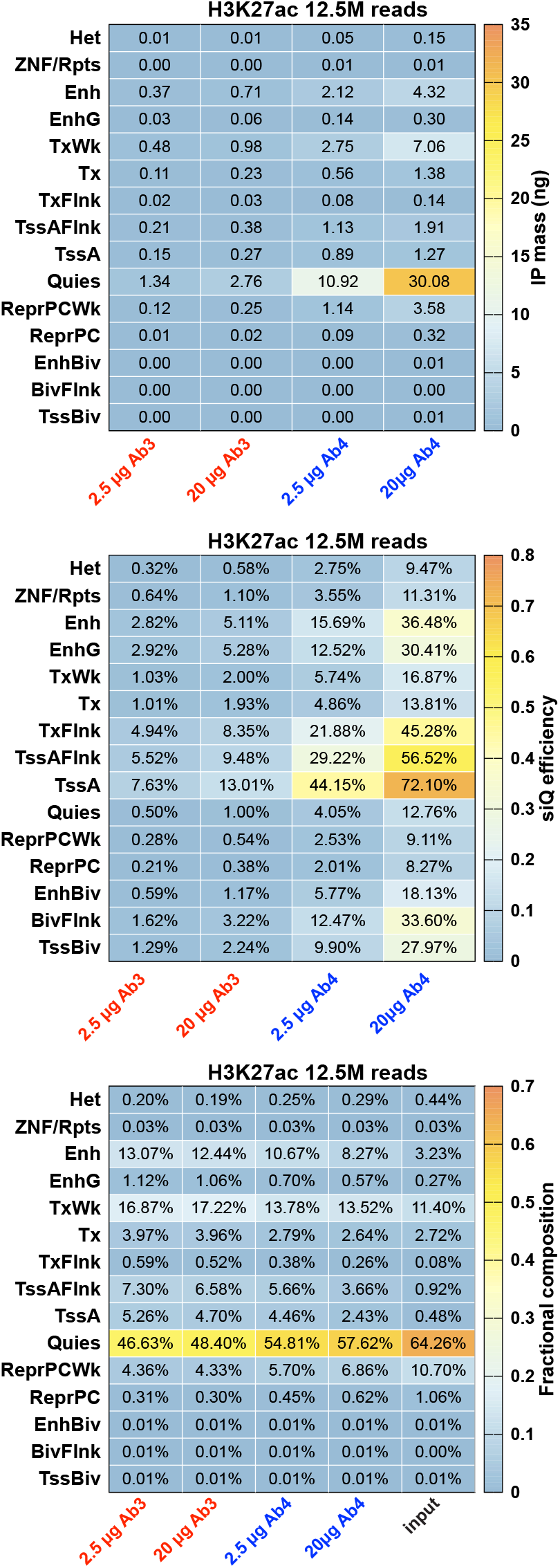
Global sequencing analysis of H3K27ac antibody titration IPs at 12.5M reads Binding isotherms of sequenced reads are labeled “12.5M” from Fig. 4A. Heatmaps of whole genome analysis of sequenced reads are shown as IP mass distribution (top), siQ-ChIP capture efficiency (middle), and fractional composition of total reads (bottom). Genome annotations are from the 15-state ChromHMM model described in reference 22.

## REFERENCES

1. Johnson, D. S., Mortazavi, A., Myers, R. M. & Wold, B. Genome-wide mapping of in vivo protein-DNA interactions. Science 316, 1497–1502 (2007).

2. Barski, A. et al. High-resolution profiling of histone methylations in the human genome. Cell 129, 823–837 (2007).

3. Dickson, B. M. et al. A physical basis for quantitative ChIP-sequencing. J Biol Chem 295, 15826–15837 (2020).

4. Fischle, W. A new approach for quantifying epigenetic landscapes. Journal of Biological Chemistry 295, 15838–15839 (2020).

5. Orlando, D. A. et al. Quantitative ChIP-Seq normalization reveals global modulation of the epigenome. Cell Rep 9, 1163–1170 (2014).

6. Grzybowski, A. T., Shah, R. N., Richter, W. F. & Ruthenburg, A. J. Native internally calibrated chromatin immunoprecipitation for quantitative studies of histone post-translational modifications. Nat Protoc 14, 3275–3302 (2019).

7. Wu, D., Wang, L. & Huang, H. Protocol to apply spike-in ChIP-seq to capture massive histone acetylation in human cells. STAR Protoc 2, 100681 (2021).

8. Chen, K. et al. The Overlooked Fact: Fundamental Need for Spike-In Control for Virtually All Genome-Wide Analyses. Mol Cell Biol 36, 662–667 (2015).

9. ENCODE Project Consortium. An integrated encyclopedia of DNA elements in the human genome. Nature 489, 57–74 (2012).

10. Dickson, B. M., Kupai, A., Vaughan, R. M. & Rothbart, S. B. Theoretical and practical refinements of sans spike-in quantitative ChIP-seq with application to p300/CBP inhibition. bioRxiv 2022.08.09.503331 (2022) doi:10.1101/2022.08.09.503331.

11. Rothbart, S. B. et al. An Interactive Database for the Assessment of Histone Antibody Specificity. Mol Cell 59, 502–511 (2015).

12. Egan, B. et al. An Alternative Approach to ChIP-Seq Normalization Enables Detection of Genome-Wide Changes in Histone H3 Lysine 27 Trimethylation upon EZH2 Inhibition. PLoS One 11, e0166438 (2016).

13. Solomon, M. J., Larsen, P. L. & Varshavsky, A. Mapping protein-DNA interactions in vivo with formaldehyde: evidence that histone H4 is retained on a highly transcribed gene. Cell 53, 937–947 (1988).

14. Wu, C.-H. et al. Sequence-specific capture of protein-DNA complexes for mass spectrometric protein identification. PLoS One 6, e26217 (2011).

15. de Jonge, W. J., Brok, M., Kemmeren, P. & Holstege, F. C. P. An Optimized Chromatin Immunoprecipitation Protocol for Quantification of Protein-DNA Interactions. STAR Protoc 1, 100020 (2020).

16. Texari, L. et al. An optimized protocol for rapid, sensitive and robust on-bead ChIP-seq from primary cells. STAR Protoc 2, 100358 (2021).

17. Zhong, L., Martinez-Pastor, B., Silberman, D. M., Sebastian, C. & Mostoslavsky, R. METHODS IN MOLECULAR BIOLOGY: ASSAYING CHROMATIN SIRTUINS. Methods Mol Biol 1077, 149–163 (2013).

18. Toyama, B. H. et al. Visualization of long-lived proteins reveals age mosaicism within nuclei of postmitotic cells. J Cell Biol 218, 433–444 (2019).

19. Arrigoni, L. et al. Standardizing chromatin research: a simple and universal method for ChIP-seq. Nucleic Acids Res 44, e67 (2016).

20. Sulkowski, E. & Laskowski, M. Mechanism of action of micrococcal nuclease on deoxyribonucleic acid. J Biol Chem 237, 2620–2625 (1962).

21. Chereji, R. V., Bryson, T. D. & Henikoff, S. Quantitative MNase-seq accurately maps nucleosome occupancy levels. Genome Biol 20, 198 (2019).

22. Ernst, J. & Kellis, M. ChromHMM: automating chromatin state discovery and characterization. Nat Methods 9, 215–216 (2012).

23. Roadmap Epigenomics Consortium et al. Integrative analysis of 111 reference human epigenomes. Nature 518, 317–330 (2015).

24. Zhang, J. et al. An integrative ENCODE resource for cancer genomics. Nat Commun 11, 3696 (2020).

25. Leroy, G. et al. A quantitative atlas of histone modification signatures from human cancer cells. Epigenetics Chromatin 6, 20 (2013).

26. Ernst, J. et al. Systematic analysis of chromatin state dynamics in nine human cell types. Nature 473, 43–49 (2011).

27. Kimura, H. Histone modifications for human epigenome analysis. J Hum Genet 58, 439–445 (2013).

28. Creyghton, M. P. et al. Histone H3K27ac separates active from poised enhancers and predicts developmental state. Proceedings of the National Academy of Sciences 107, 21931–21936 (2010).

29. Zhang, T., Zhang, Z., Dong, Q., Xiong, J. & Zhu, B. Histone H3K27 acetylation is dispensable for enhancer activity in mouse embryonic stem cells. Genome Biology 21, 45 (2020).

30. Zhong, J. et al. Purification of nanogram-range immunoprecipitated DNA in ChIP-seq application. BMC Genomics 18, 985 (2017).

31. Edgar, R., Domrachev, M. & Lash, A. E. Gene Expression Omnibus: NCBI gene expression and hybridization array data repository. Nucleic Acids Res 30, 207–210 (2002).

32. Cornett, E. M., Dickson, B. M. & Rothbart, S. B. Analysis of Histone Antibody Specificity with Peptide Microarrays. JoVE (Journal of Visualized Experiments) e55912 (2017) doi:10.3791/55912.

33. Dickson, B. M., Cornett, E. M., Ramjan, Z. & Rothbart, S. B. ArrayNinja: An Open Source Platform for Unified Planning and Analysis of Microarray Experiments. Methods Enzymol 574, 53–77 (2016).

